# Bio-upcycling of polyethylene terephthalate

**DOI:** 10.1101/2020.03.16.993592

**Authors:** Till Tiso, Tanja Narancic, Ren Wei, Eric Pollet, Niall Beagan, Katja Schröder, Annett Honak, Mengying Jiang, Shane T. Kenny, Nick Wierckx, Rémi Perrin, Luc Avérous, Wolfgang Zimmermann, Kevin O’Connor, Lars M. Blank

## Abstract

Over 359 million tons of plastics were produced worldwide in 2018, with significant growth expected in the near future, resulting in the global challenge of end-of-life management. The recent identification of enzymes that degrade plastics previously considered non-biodegradable opens up opportunities to steer the plastic recycling industry into the realm of biotechnology. Here, we present the sequential conversion of polyethylene terephthalate (PET) into two types of bioplastics: a medium chain-length polyhydroxyalkanoate (PHA) and a novel bio-based poly(amide urethane) (bio-PU). PET films were hydrolyzed by a thermostable polyester hydrolase yielding 100% terephthalate and ethylene glycol. A terephthalate-degrading *Pseudomonas* was evolved to also metabolize ethylene glycol and subsequently produced PHA. The strain was further modified to secrete hydroxyalkanoyloxy-alkanoates (HAAs), which were used as monomers for the chemo-catalytic synthesis of bio-PU. In short, we present a novel value-chain for PET upcycling, adding technological flexibility to the global challenge of end-of-life management of plastics.

**Graphical abstract:** 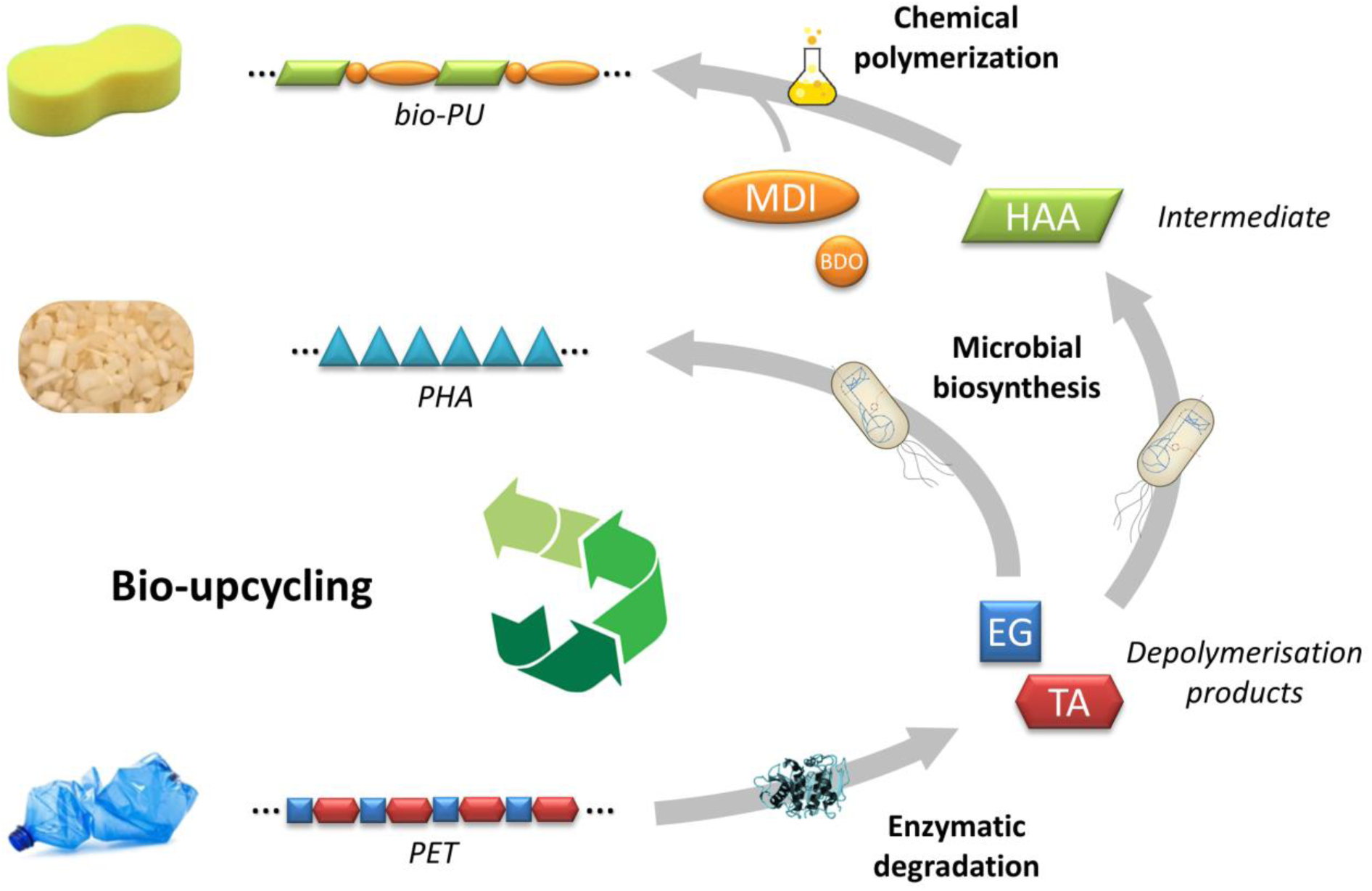

## 1 Introduction

One of the challenges humankind faces is the shift to a sustainable plastic industry. In 2018, 359 million tons of plastics have been produced worldwide and this number is growing at a rate of approximately 3% per *annum*^*1*^. Of all the plastic ever produced, only 9% was recycled and 12% was incinerated. The remaining majority is either in use or was landfilled, with a chance to be released into the environment^2^. Indeed, in 2010 an estimated 5-13 million tons of plastic ultimately ended up in the ocean^3^. While plastic, due to its lightweight and sturdiness, has many environmentally beneficial applications, the environmental damage caused by plastic must be arrested by addressing the end-of-life challenge.

State-of-the art plastic recycling is either via mechanical or chemical methods, or a combination thereof^4^. An ideal plastic for recycling is polyethylene terephthalate (PET). The main PET product, beverage bottles, can be specifically collected, avoiding mixed material challenges. In addition, with its thermoplastic properties such as high melting temperature and the possibility to process it without the use of additives, PET fulfils many technical recycling criteria. While in some European countries, PET is collected at quotas above 95%, only approximately 30% of it is recycled, even under these ideal conditions^5^. Reasons are manifold including cost, consumer acceptance, and safety regulations surrounding recycled material, to name a few. An alternative way to increase plastic recycling is to add additional value to the plastic waste, not aiming for the same material or consumer good (e.g., bottle-to-bottle recycling), but rather upcycling to chemicals and materials of higher value. This concept has already been demonstrated using chemical methods as glycolization^6^, alcoholysis^7^, glycolysis^8,9^, and organocatalysis^10^.

This upcycling can potentially be achieved by using carbon-rich plastic waste streams as a substrate for biotechnological processes^5^. Here, PET is degraded into its monomers terephthalic acid and ethylene glycol and used as carbon and energy feedstock for microbes that produce valuable molecules and materials.

In 2014 Yang *et al*.^11^ reported that the larvae of the meal moth *Plodia interpunctella* can degrade polyethylene, a trait also discovered later in related species^12^. Two bacterial species from the gut of this meal moth larvae were likely responsible the degradation of polyethylene^11^. Similarly, in 2015, *Exiguobacterium* was identified as a polystyrene degrading organism^13^ from polystyrene-eating mealworms^14^. In 2016 the bacterium *Ideonella sakaiensis* was reported to degrade amorphous PET when cultured in the presence of yeast extract as an additional carbon source^15^. The molecular basis of the ester-bond hydrolyzing PETase and mono-(2-hydroxyethyl)TA (MHET)ase enzymes of this strain was reported in several publications (e.g.,^16,17^).

For obvious reasons, the biodegradation of these recalcitrant plastics are exciting discoveries that give hope for the natural bioremediation of sites contaminated with plastic waste in the environment, although plastic degradation in the ocean seems to be slow at best and the anthropogenic dissemination of new plastic pollution likely far exceeds its decay^18^. Notably, this biodegradation also offers a tremendous opportunity for waste treatment: To biotechnologically upcycle plastic waste to valuable products such as bioplastics. In principle, as we see it^5,19^, plastic waste biotechnology mirrors the well-known utilization of lignocellulosic hydrolysate: i) Enzymatic hydrolysis of the polymeric substrate, ii) metabolism of the resulting hydrolysates by microorganisms, and iii) production of value-added chemicals and polymers by these organisms. However, unlike plant biomass, plastics are often chemically less complex consisting of only a few well-defined monomers, making them potentially much easier substrates for biotechnological utilization. PET, for instance, is a highly pure polymer compared to biomass, composed of almost 100% ethylene glycol (EG) and terephthalic acid (TA)^20,21^. The majority of commercially used PET is a semi-crystalline polymer with a significant amorphous content, which is particularly amenable to enzymatic depolymerization at its glass transition temperature of 70 °C^22^. Therefore, compared to the mesophilic *I. sakaiensis* enzymes, counterparts from thermophilic microorganisms stable at > 70 °C emerged as more promising biocatalysts for the rapid degradation of PET plastic waste^20,23^. However, the microbial degradation of lignocellulose, which is biotechnologically challenging by itself^24^, has an evolutionary head start of hundreds of millions of years, considering that plastics have only entered the biosphere in the last century. The need for optimization of novel biotechnological catalysts thus becomes obvious. Enzymes, microbes, and processes are required that are capable of degrading synthetic polymers previously considered as non-biodegradable with high rates.

Here, we present the biocatalytic upcycling of PET into two types of biopolymers using a multidisciplinary approach (Figure 1). PET was hydrolyzed enzymatically in a dedicated reactor into its monomers EG and TA. The monomers were converted by a modified *Pseudomonas* sp. GO16 into the native intracellular polymer polyhydroxyalkanoate (PHA) and into the engineered extracellular building block hydroxyalkanoyloxy-alkanoate (HAA). After HAA purification, this platform molecule^25^ was chemically co-polymerized to form a novel partly bio-based poly(amide urethane) (bio-PU). In short, we present novel sequential bio-upcycling routes for PET waste, adding technological flexibility to the global challenge of sustainable end-of-life management of plastics.

**Figure 1:**
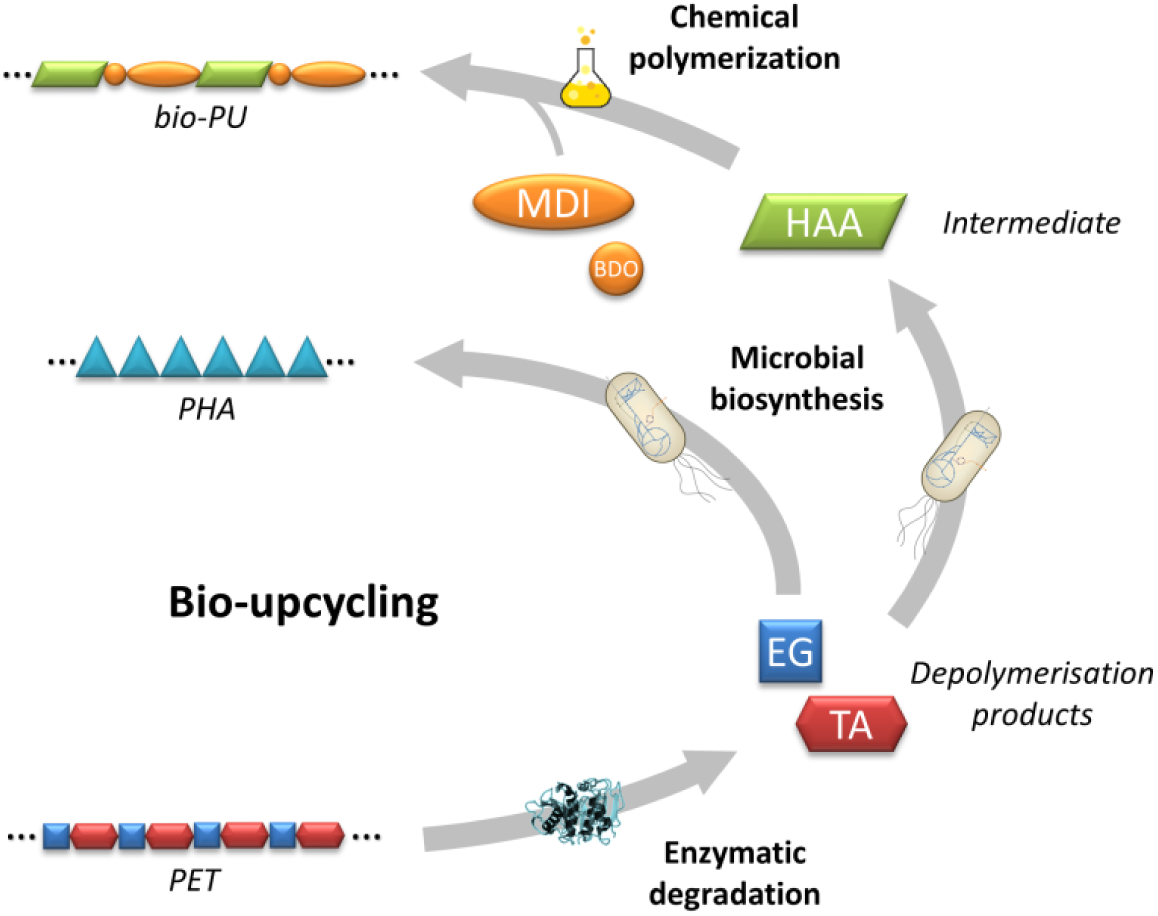
Two alternative bio-upcycling routes for PET waste to novel biopolymers. PET hydrolysis is catalyzed by a thermostable polyester hydrolase. The resulting monomers EG and TA (blue and red, respectively) are fed to an engineered *P. putida*, which synthesizes either extracellular HAA (green), or the intracellular biopolymer PHA (cyan). While PHA is already a polymer, HAA is co-polymerized with a diisocyanate and butanediol (orange) to yield a novel bio-based poly(amide urethane) (bio-PU).

## 2 Results

### 2.1 Enzymatic PET degradation

The here presented interdisciplinary approach for the upcycling of PET is initiated by enzymatic cleavage of the polymer. Microbial polyester hydrolases capable of efficiently degrading amorphous or low-crystallinity PET samples^26^ at elevated temperatures close to the glass-transition temperature have been found in fungi^27^, thermophilic actinomycetes (e.g.,^20,28-30^), and in plant compost^31^. Leaf-branch compost cutinase (LCC) is a polyester hydrolase, of which the encoding gene was originally isolated from a plant compost metagenome^31^. The enzyme was produced in *E. coli* and purified as described previously^32^.

A single scaled-up PET hydrolysis experiment was carried out to generate the material for the subsequent steps. Figure 2 shows the time courses of the formation of the degradation products released from amorphous PET films during LCC-catalyzed hydrolysis in a stirred tank reactor (STR), as well as of the residual esterolytic activity and pH values determined in the reaction supernatant. The concentration of TA and EG showed a steep near-linear increase within the first 24 h of the hydrolytic reaction and weakened to a markedly lower rate until 120 h. By contrast, the amount of MHET increased gradually in the early stage of the reaction and peaked at 48 h, followed by a decline until its complete disappearance at 96 h. Afterwards, TA was the only detectable soluble UV-absorbing degradation product in the supernatant. In previous studies, MHET was found to be an inhibitor for polyester hydrolases including LCC (e.g.,^33,34^). Compared to the ester bonds in PET, this mono-ester of TA and EG was not preferentially cleaved. Therefore, LCC was assumed to preferably catalyze the depolymerization of PET during the first 48 h, followed by a detectable cleavage of MHET until its complete disappearance in the reaction supernatant after 96 h. According to the chemical structure of PET, EG and TA should have been released in equivalent amounts. However, a slightly higher amount of EG was detected from 24 h onwards, presumably due to the fact that a small excess fraction of EG can always be found as dimers (DEG) in PET polymers as a result of synthesis^20^. After 120 h, 215.6 mM TA and 250.4 mM EG were detected in the reaction supernatant (Figure 2). The rapid release of TA in the first 24 h caused a continuous pH decrease, even though a high buffer concentration of 1 M potassium phosphate was used. As the TA formation rate decreased after 72 h, the pH value determined in the reaction supernatant remained comparably stable. Extending the reaction time to 168 h, the PET film (15.7 g) was almost completely (nearly 100%) broken down into soluble low molecular weight compounds, leaving only several tiny fibers with neglectable weight (<0.5 mg). The esterolytic activity indicates a thermal activation of LCC in the first 8 h, followed by a sharp decline back to its initial value. Afterwards it remained almost unchanged until 120 h, indicating a high thermal stability of LCC at 70 °C, which is in a good agreement with a previous study^35^.

**Figure 2:**
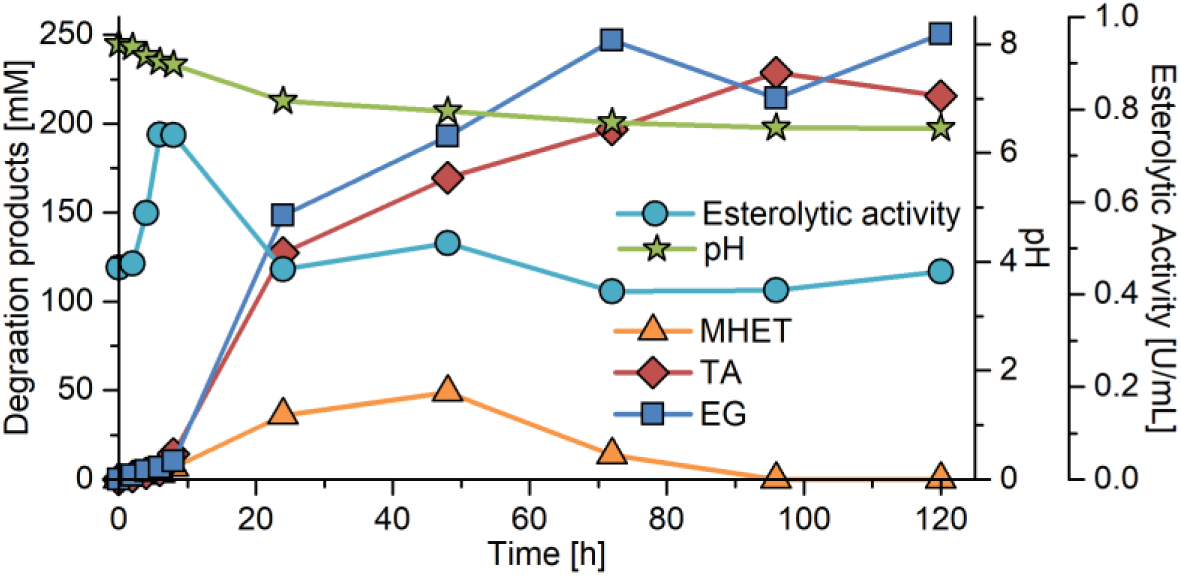
PET film hydrolysis. Time courses of PET hydrolysis into (soluble) ethylene glycol (EG), terephthalic acid (TA), and mono-(2-hydroxyethyl)TA (MHET) (primary y-axis) catalyzed by purified LCC in a stirred tank reactor at 70 °C, as well as of pH value and residual esterolytic activity in the reaction supernatant (secondary/tertiary y-axis). Data from a single representative experiment.

### 2.2 Upcycling of enzymatically hydrolyzed PET

Two different pathways were employed for the biosynthesis of molecules for bioplastic production using *Pseudomonas*. Pseudomonads are Gram-negative bacteria with a high potential for degrading synthetic plastics^5,36^, due to their versatile arsenal of catabolic enzymes^37^. Different *Pseudomonas putida* strains are known for their metabolism of a wide variety of substrates, including aromatics such as TA^38^, and aliphatics such as EG^39-41^. Indeed *Pseudomonas* species strain GO16 (isolated from a PET bottle processing plant) is able to metabolize TA and was previously used to produce the biodegradable polymer PHA from TA emanating from pyrolyzed PET^38^.

#### 2.2.1 Growth of *Pseudomonas* sp. GO16 on PET monomers

To enable conversion of both monomers of PET, metabolization of EG had to be established. It was observed that *Pseudomonas* sp. GO16 was capable of slow growth on EG with a lag phase of more than 5 days. This growth was improved by adaptive laboratory evolution as also shown previously for *P. putida* KT2440^42^. After more than 45 days of repeated batch cultivations, *Pseudomonas* sp. GO16 KS3 was able to grow on EG at a rate of 0.4 h^-1^ (data not shown).

#### 2.2.2 Synthesis of the biopolymer polyhydroxyalkanoate (PHA)

The monomers from PET hydrolysis were used for conversion into PHA using *Pseudomonas* sp. GO16 KS3. PHAs are a family of bacterial carbon and energy storage polyesters which represented 2% of the global bioplastic market in 2014^43^. With over 150 known PHA monomers, (*R*)-3-hydroxyalkanoic acids, PHAs have highly diverse material properties and therefore a broad range of applications from packaging to medical^44^.

The conversion of hydrolyzed PET (described above) into PHA was carried out in a 5 L batch reactor. TA was completely consumed by 23 h of cultivation, while EG was consumed at a 3.5-fold lower rate and did not reach complete exhaustion (Figure 3). When using a synthetic mixture of TA and EG, the EG depletion rate was much higher (see supplementary information). The highest biomass achieved was 2.3 g/L after 22 h of incubation, remaining at a similar level for the next 5 h. Nitrogen was completely exhausted after 16 h of cultivation, corresponding to the onset of PHA accumulation. The PHA level kept rising after TA depletion, and reached a maximum of 0.15 g/L representing approximately 7% of cell dry weight (CDW) and indicating that both TA and EG derived from enzymatically hydrolyzed PET were converted into PHA (Figure 3). The biomass yield, including PHA, was 0.4 g/g. The medium-chain-length PHA produced by *Pseudomonas* sp. GO16 KS3 from hydrolyzed PET consisted of C_10_ (61 mol%), C_12_ (24 mol%), and C_8_ (15 mol%).

**Figure 3:**
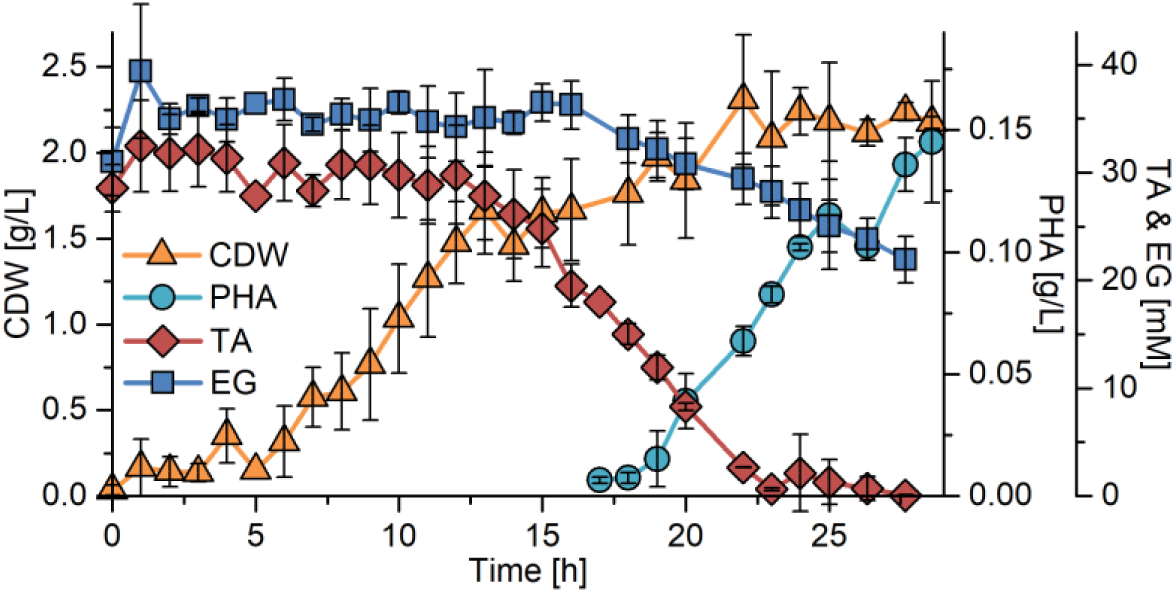
Growth, PHA accumulation, and substrate depletion by *Pseudomonas* sp. GO16 KS3 when enzymatically hydrolyzed PET was used under nitrogen limiting conditions. *Pseudomonas* sp. GO16 KS3 was cultivated in a 5 L bioreactor with 3 L of mineral salts medium (MSM) at 30 °C. The hydrolyzed PET was added to a concentration of approximately 30 mM TA and EG. Growth (cell dry weight, CDW), polyhydroxyalkanoate (PHA, %CDW), substrates terephthalic acid (TA) and ethylene glycol (EG). The error bars represent the standard deviation from the mean of three independent biological replicates.

#### 2.2.3 Biosynthesis of extracellular HAA

While many *Pseudomonas* strains are known to natively produce PHAs, pseudomonads have further been engineered to produce a broad palette of other industrially relevant platform chemicals^37,45,46^. Recently, *P. putida* KT2440 was engineered to synthesize HAA, a dimer of 3-hydroxy fatty acids. In contrast to many other fatty acid containing molecules, HAAs are secreted into the growth medium^47^, resulting in simpler purification that does not rely on cell lysis. HAAs are amphiphilic molecules with surface-active properties and interesting platform chemicals for further bio- or chemo-catalytic conversion^25^.

Thus, in order to demonstrate the production of a non-native value-added molecule from hydrolyzed PET, the evolved *Pseudomonas* sp. GO16 KS3, capable of growth with TA and EG, was transformed with the HAA synthesis plasmid pSB01^47^. The PET hydrolysis solution was diluted to meet the requirements for the microbe and essential nutrients (nitrogen source, trace elements) were added. The resulting concentrations of EG and TA were approximately 15-18 mmol/L.

With this medium, an HAA concentration of 35 mg/L was achieved (Figure 4a). During the first ten hours, the TA concentration declined rapidly. EG was only taken up after TA was completely consumed, taking approximately 10 additional hours. However, the engineered strain only synthesized HAA from TA, as HAA reached maximum concentration after 12 h, corresponding to TA depletion. EG apparently was used for growth only, since the cell dry weight increases further while EG is consumed after a short transition phase. The production rate amounted to 5 mg/L/h while the yield was 0.01 g_HAA_/g_TA_. The theoretical yield for HAA synthesis from TA is 0.39 g_HAA_/g_TA_ as determined by flux balance analysis.

**Figure 4:**
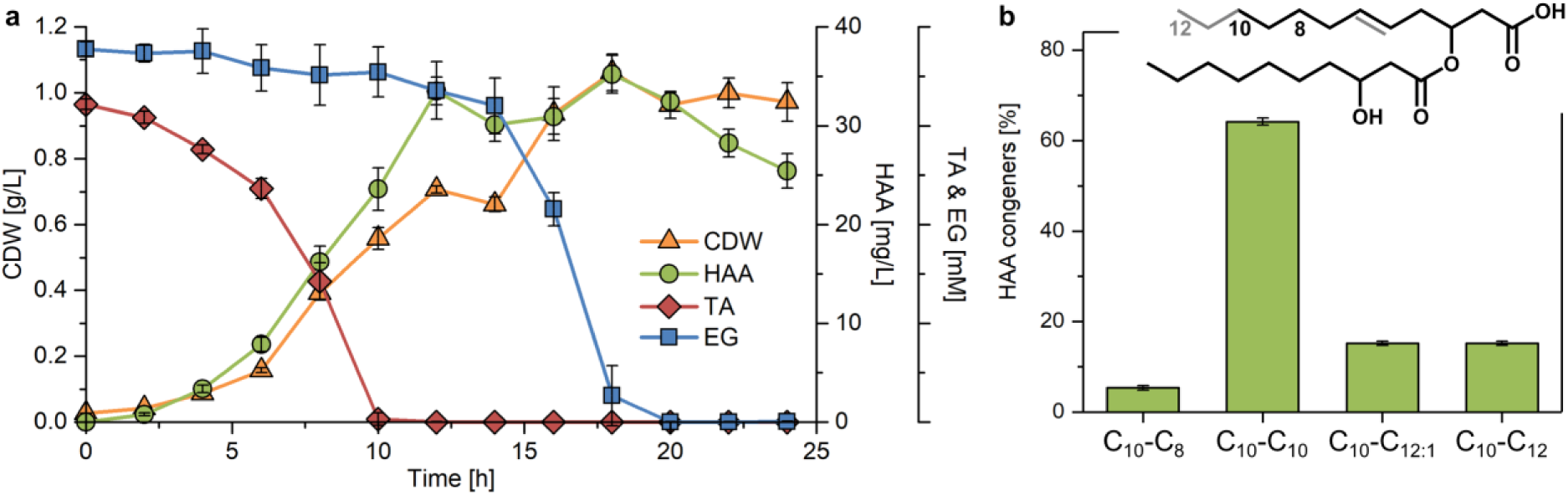
**a) HAA synthesis from hydrolyzed PET.** *Pseudomonas* sp. GO16 KS3 pSB01 was cultivated in shake flasks with 50 mL of mineral salts medium (MSM) at 30 °C. The PET hydrolysate was added to a concentration of 15-18 mM TA and EG. Growth (cell dry weight, CDW), HAA, substrates terephthalic acid (TA) and ethylene glycol (EG) depletion. The error bars represent the deviation from the mean of two biological replicates. **b) Molecular diversity of the HAA congeners synthesized from PET monomers.** The error bars represent the standard deviation from the mean of two independent fermentations (as indicated in a) at four time points, i.e. eight biological replicates.

The engineered *P. putida* synthesizes a mixture of four HAA congeners as identified by HPLC-CAD (Figure 4b). The mainly produced hydroxy fatty acid detected was hydroxydecanoate. The length of the second monomer varied between eight and twelve carbon atoms, of which the C_12_ can be unsaturated.

### 2.3 Polymerization of HAA

The final stage of the process is the chemical polymerization of the biotechnologically produced HAA to produce bio-PU. Since an isocyanate moiety can react with both an hydroxyl and a carboxylic acid group, and HAA is an hydroxy acid, its direct polymerization with 4,4’-methylene diphenyl diisocyanate (MDI) and butanediol (BDO) was performed and led to the formation of a poly(amide urethane). This polymer is still partly based on petrochemical materials. While bio-BDO is available^48^, bio-anilin patents^49^ suggest that bio-MDI will be available in the future^50^, to render the synthesized PU completely bio-based. The length of the HAA side chain can be varied depending on the RhlA used for synthesis^51^, thereby influencing plastic properties.

#### 2.3.1 Direct polymerization to poly(amide urethane)

The direct polymerization of HAA with MDI has been carried out by a two-step polymerization involving the synthesis of a prepolymer followed by a chain extension reaction with BDO (Figure 5).

**Figure 5:**
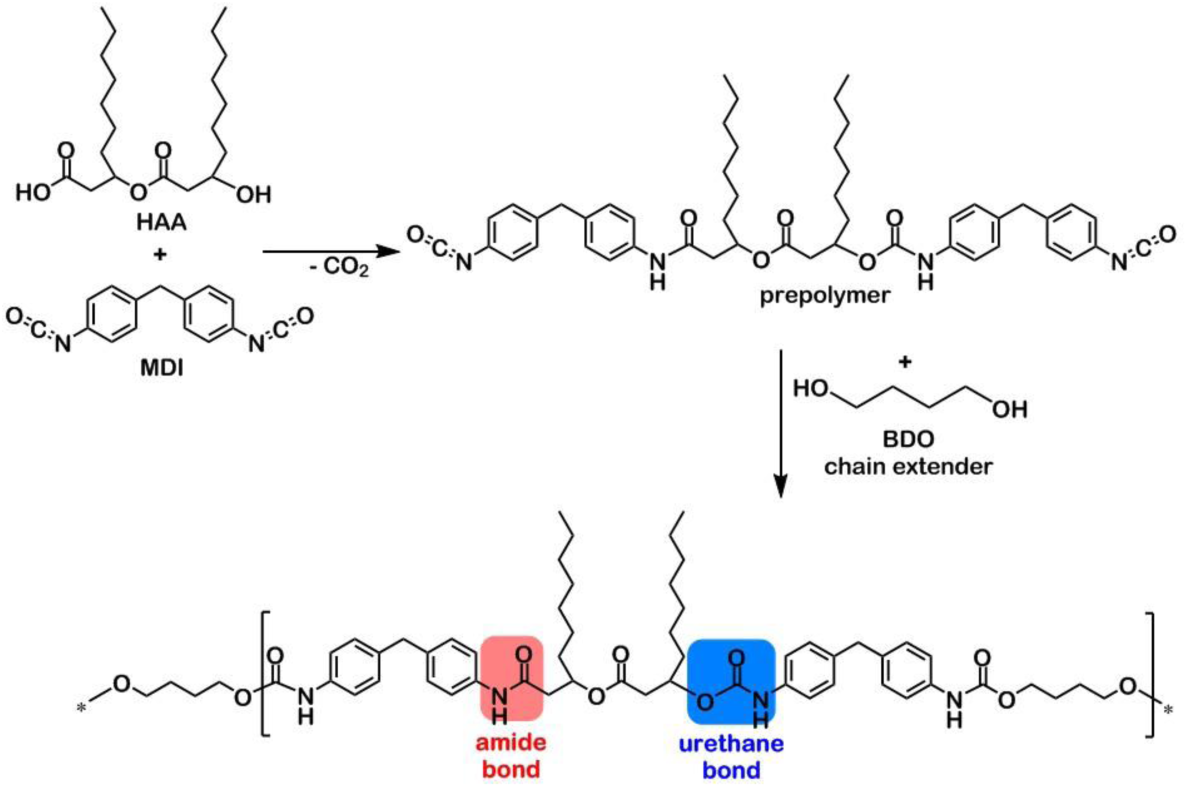
Novel bio-PU synthesis. Polymerization reaction between the hydroxy fatty acid ester HAA, and MDI diisocyanate. The resulting prepolymer is submitted to chain extension with BDO to form a partly bio-based poly(amide urethane) (bio-PU).

FTIR and NMR analyses (Figure 6) indicated simultaneous amide and urethane bond formation, while the reactivity of 4,4’-MDI with the secondary alcohol appeared to be slightly higher than with the carboxylic acid. These analyses confirmed the synthesis of a novel poly(amide urethane).

**Figure 6:**
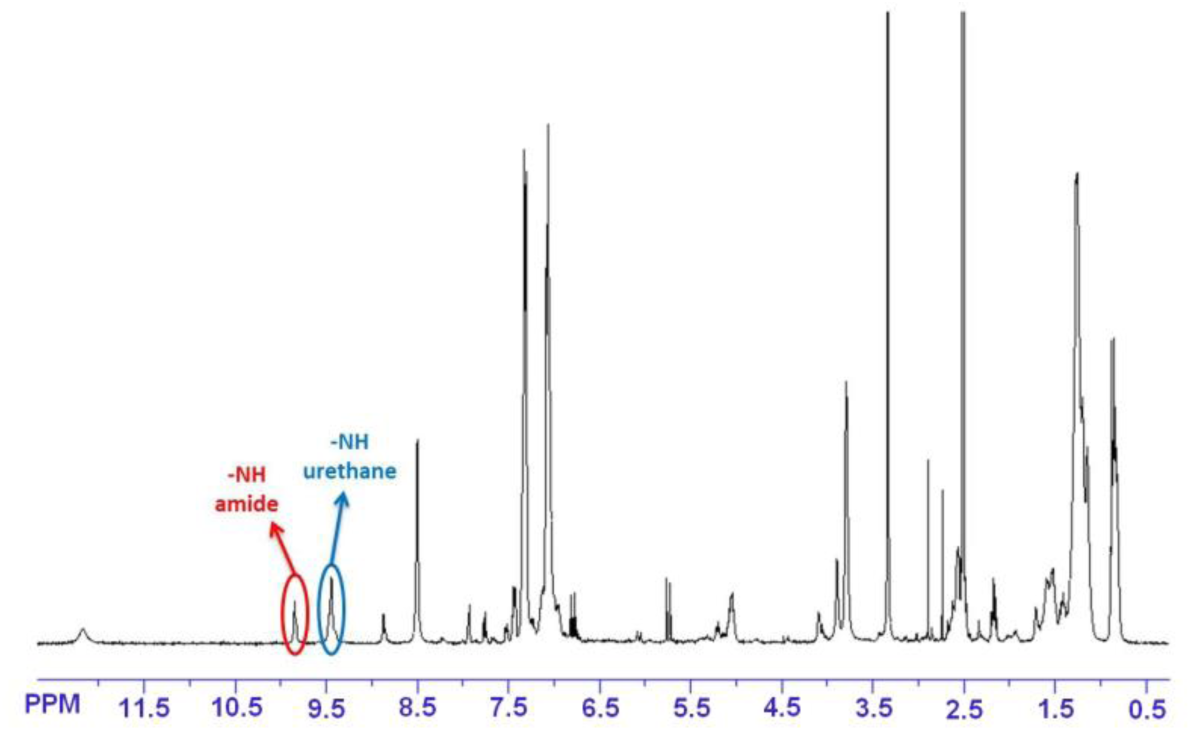
Confirmation of bio-PU synthesis. 1H NMR spectrum of the final polymer showing the presence of NH signals of both amide (circled in red) and urethane (circled in blue) bonds.

Differential scanning calorimetry (DSC) analysis of the bio-PU evidenced the amorphous character of the polymer and revealed a glass-transition temperature of 50 °C. This value is consistent with the glass-transition temperature usually reported for polyurethanes obtained from MDI and BDO as main diol, which may vary from 60 to 110 °C^52^. The slightly lower value obtained here can be explained by the HAA displaying two pendant groups once polymerized with MDI (Figure 5). These aliphatic pendant groups may increase the chain mobility and can thus be used to tune down the glass-transition temperature in comparison with polyurethane only made from BDO as diol.

The thermal stability of this HAA-based polyurethane was determined by thermogravimetric analysis (TGA). The polymer started to degrade and to lose volatile products at 160 °C and then showed a multi-step degradation profile with the main mass loss occurring between 250 and 350 °C. This behavior is consistent with the thermal stability usually observed for such polymer systems. The rather low onset degradation temperature could be explained by the presence of the ester bond in the HAA, which makes it more sensitive to thermal degradation (the temperature of the maximum degradation rate of HAA is around 215 °C).

## 3 Discussion

We present the bio-upcycling of previously considered non-biodegradable plastic waste by cascading the enzymatic depolymerization of PET with the microbial conversion into valuable polymers. Renewable plastics including PHA have already been proposed to effectuate a shift of the packaging industry, which consumes over 38% of the plastics produced^53^. The results exemplify the previously proposed value-chain for the utilization of plastic waste as additional carbon source in biotechnology to produce a wide range of valuable products^5,19^. We see great potential in this new approach to recycling, and thus consider this study a starting point for new research in enzyme technology, strain engineering, and polymer chemistry, akin to the megatrends in lignocellulosic biotechnology that we have seen in the last decades^54,55^.

Lignocellulosic biotechnology has already found solutions to the challenges we are about to face in plastic waste biotechnology. Parameters like high solid loads and enzyme amounts have to be optimized using heuristic approaches and trial-and-error methodologies. The use of enzyme engineering and enzyme cocktail formulation will enable an even more efficient PET degradation, for instance using specialized enzymes of the various types of PET, i.e. high molecular weight PET and PET oligomers mono-(2-hydroxyethyl)TA (MHET) and bis-2(hydroxyethyl)TA (BHET); possibly combined with chemical hydrolysis methods such as glycolysis^56^. The use of enzyme cocktails will also enable feedstock flexibility, especially when combined with microbes engineered to accept other plastic-derived substrates. For example, the novel PET-like polymer polyethylene furanoate (PEF) is also degraded by a PET hydrolase^17^, and *P. putida* can be engineered to degrade the resulting 2,5-furandicarboxylic acid^57,58^. It is worth noting that logistics is a major hurdle in lignocellulosic biotech since often completely new infrastructure has to be built (*i.e*., from forest to factory). In contrast, many countries already have highly efficient plastic waste collection systems, which can find a new valorization through the herein proposed bio-upcycling route.

To achieve the latter will require significant research and technology development. Learning again from lignocellulosic biotech, enzyme discovery and engineering is key for resource- and cost-efficient hydrolysis of the polymers. For PET, the abundance of polyester hydrolases in the environment was investigated in detail. The results suggest that this enzyme activity is rather rare, but more frequent in crude oil rich environments^26,59,60^. Currently further metagenome and mechanistic studies of this important enzyme class are carried out by the scientific community, most likely discovering protein family members with superb activities or at least interesting amino acid variations. The latter can be exploited in protein engineering efforts, already published for the PETase of *I. sakaiensis*^61^ and thermophilic proteins with the same enzyme specificity^17,22,62^.

We here show the complete degradation of PET, enabling a range of process optimization strategies, which is a significant advantage over lignocellulose-based processes. Lignocellulose-derived substrates come with a large fraction of solids, which are not completely degraded impeding the application of, e.g., enzyme immobilization or *in situ* removal of formed monomers (by e.g., precipitation or extraction).

While the mesophilic PET hydrolase from *I. sakaiensis*^15^ suggests consolidated hydrolysis and utilization, we focused on sequential plastic depolymerization and monomer conversion on purpose. This approach provides a higher flexibility for optimizing of process conditions and (bio)catalysts used. Another advantage is the built-in pasteurization step (PET hydrolysis at 70 °C), which renders the resulting solution of monomers from PET hydrolysis semi sterile. If the feedstock is for example food package waste, sterility is a huge problem.

Accessing non-biodegradable plastics of petrochemical origin (and in the future of biological origin) as carbon source for fermentations enables biotechnology to valorize enormous waste streams for the sustainable production of many valuable products by exploiting the metabolic versatility of microorganisms. Products such as aromatics, organic acids, glycolipids and lipid derivatives as well as biopolymers and fuel molecules are just some examples^46,47,63,64^. More importantly, the established biosynthetic pathways for plastic monomer metabolism can also be adapted to function in different organisms, widening the applicability of the described approach even further. The production of novel bioplastics moreover entails the advantage that resulting products can be bio-upcycled more efficiently using the described approach as they often feature a higher biodegradability. Plastic waste biotechnological upcycling thus offers novel possibilities of end-of-life management by closing multi-million-ton material cycles in a circular economy, tackling two challenges of our petrochemical economy at the same time: Arresting the unrestrained consumption of crude oil and the resulting emission of greenhouse gases, as well as the pollution of our environment with plastic waste.

## 4 Methods

### 4.1 Chemicals

Suprasec 2385 (uretonimine-modified 4,4’-diphenylmethylene diisocyanate (4,4’-MDI)) was supplied by Huntsman. 1,4-butanediol (1,4-BDO) (99%), N,N-dimethylformamide (DMF), ethyl acetate, pyridine, p-toluenesulfonic acid (p-TSA), oxalyl chloride (98%), deuterated chloroform (CDCl_3_) and deuterated dimethyl sulfoxide (DMSO-d_6_) were purchased from Sigma-Aldrich. All solvents used for the analytical methods were of analytical grade.

### 4.2 Bacterial strains and plasmids

Strain *Pseudomonas* sp. GO16 (accession number NCIMB 41538, NCIMB Aberdeen, Scotland, UK) was used for PHA and HAA synthesis. *Escherichia coli* BL21 (DE3) was used to produce the recombinant polyester hydrolase LC-cutinase^31^. *Pseudomonas* sp. GO16 KS3 was transformed with pSB01, the plasmid mediating HAA synthesis constructed previously^47^.

### 4.3 LC-cutinase production

The recombinant enzyme production was carried out in *E. coli* BL21 (DE3) harboring pET-20b(+) containing the synthetic gene^65^ (ENA: LN879395) encoding LCC, which was originally identified from a plant compost metagenome. A 42-L fermenter (Infors AG, Bottmingen, Switzerland) with a working volume of 25 L was used to produce recombinant LCC as described previously for a homologous polyester hydrolase^66^.

Briefly, the recombinant *E. coli* culture was grown at 37 °C to an optical density at 600 nm (OD_600_) of 1.5. The recombinant protein production was induced at a final IPTG concentration of 0.5 mM at 18 °C for 14 h. Bacterial cells were harvested by centrifugation at 11,285×g and 4 °C for 25 min. Cell pellets were resuspended in 50 mM sodium phosphate buffer (pH 8) containing 300 mM NaCl and disrupted by ultra-sonication. Following the removal of cell debris by centrifugation, the resulting supernatant containing soluble LCC was subjected to purification by immobilized metal ion chromatography (IMAC) using Ni-NTA Superflow (Qiagen, Hilden, Germany). Protein elution using 250 mM imidazole was then dialyzed against 100 mM potassium phosphate buffer (pH 8) prior to the application in the enzymatic PET hydrolysis. Protein content was determined using the Bradford method. The esterolytic activity was determined using *para*-nitrophenyl butyrate (*p*-NPB) as a substrate as described elsewhere^67^. One unit esterolytic activity was defined as the amount of enzyme required to hydrolyze 1 µmol of *p*-NPB per min.

### 4.4 Enzymatic hydrolysis of PET

The enzymatic hydrolysis of PET was carried out in a 1 L temperature controlled stirred tank reactor (STR, Duran Group GmbH, Wertheim/Main, Germany). Approximately 15.7 g of amorphous PET film (surface of 1,000 cm^2^) (product number ES301445, Goodfellow Cambridge Ltd., Huntingdon, UK) were cut into pieces of about 2×2 cm^2^, and then washed with 0.1% SDS, ethanol, and ultra-pure water, followed by drying at 50 °C for 24 h. Cleaned PET film pieces were placed in the STR containing 400 mL of 20 µg/mL IMAC purified LCC dissolved in 1 M potassium phosphate buffer (pH 8). The hydrolytic reaction was conducted at a constant temperature of 70 °C and an agitation speed of 100 rpm. Samples of 4 mL were removed from the reaction supernatant at time points 0, 2, 4, 6, 8, 24, 48, 72, 96, 120 h for offline analytics. The amounts of terephthalic acid (TA) and ethylene glycol (EG), and their mono-ester (MHET) released from the PET bulk polymer as well as the residual esterolytic activity against *p*-NPB and the pH in the reaction supernatant were determined. The enzymatic PET hydrolysis was terminated after 168 h. The resulting soluble reaction supernatant obtained by filtration using a paper filter was transferred to the microbial fermentation.

### 4.5 PHA production

*Pseudomonas* sp. GO16 KS3 was inoculated from a glycerol stock onto mineral salts medium (MSM) solidified with 1.5% agar supplemented with 4.4 g/L (20 mM) disodium terephthalate (TA; Sigma). MSM contained 9 g/L Na_2_HPO_4_·12H_2_O, 1.5 g/L KH_2_PO_4_, and 1 g/L (MSM_full_) or 0.25 g/L (MSM_lim_) NH_4_Cl. Prior to inoculation MSM was supplemented with MgSO_4_ (200 mg ml^-1^) and trace elements (per liter: 4 g ZnSO_4_·7H_2_O; 1 g MnCl·4H_2_O; 0.2 g Na_2_B_4_O_7_·10H_2_O; 0.3 g NiCl_2_·6H_2_O; 1 g Na_2_MoO_4_·2H_2_O; 1 g CuCl·2H_2_O; 7.6 g FeSO_4_·7H_2_O). A single colony was inoculated into 3 ml of MSM_Full_ supplemented with 20 mM TA and incubated for 18 h at 200 rpm and 30°C. The seed cultures for the bioreactor experiments were prepared in 50 ml MSM_full_ supplemented with a synthetic mixture of 3.32 g/L (20 mM TA) and 1.24 g/L (20 mM) EG, cultivated for 18 h at 200 rpm and 30°C.

*Pseudomonas* sp. GO16 KS3 was cultivated in a 5 L stirred tank bioreactor (Sartorius UniVessel Glass 5 l) containing 3 L of MSM_lim_ broth. The bioreactor was inoculated with 150 ml of seed culture. Each cultivation was run for 28 h at constant temperature of 30°C. Air was sparged at a rate of 3 L/min and a Rushton impeller was utilized for each experiment with a minimum stirring rate of 500 rpm. All parameters were automatically controlled to maintain a minimum dissolved oxygen (DO) level above 20% and a pH of 7.0 ± 0.1 *via* the addition of 5 M NaOH and 15% (vol/vol) H_2_SO_4_ to impose inorganic nutrient limitation and therefore stimulate PHA accumulation. The hydrolyzed PET was supplied at the amount to yield 40 mM TA and EG. Three 2 ml samples were taken at regular intervals for the analysis of TA, EG and nitrogen concentrations, biomass, and PHA accumulation for each time point. The samples were centrifuged at 16,000 *g* for 3 minutes. The supernatant was collected and stored at −20°C prior to analysis, while the cell pellet was stored at −80°C and subsequently lyophilized (Labconco FreeZone 12 bulk tray freeze drier, USA) for 24 h and weighed.

### 4.6 Adaptive laboratory evolution

As *Pseudomonas* sp. GO16 was not able to metabolize EG, adaptive laboratory evolution was carried out with the constant selective pressure of EG as the sole carbon source. This experiment was carried out in shake flasks as repetitive batch culture in duplicates and continued for almost 50 days.

### 4.7 Production of HAA

The medium for HAA synthesis was based on the mineral salt medium by Hartmans *et al.*^68^. The monomer solution from PET hydrolysis was diluted 1:20 and autoclaved. This solution was used to prepare the medium instead of water. HAA was produced and purified as shown previously^47^.

### 4.8 Direct HAA polymerization to poly(amide urethane)

The direct HAA polymerization to poly(amide urethane) was performed with an isocyanate to hydroxyl and acid molar ratio ([NCO] / ([OH] + [COOH])) equal to 2 and without catalyst addition. In a round bottom flask of 50 mL, the appropriate amount of HAA (505 mg) and modified 4,4’-MDI (787 mg) were introduced under nitrogen flux. The reaction mixture was heated up to 90 °C and magnetically stirred for 4 h under nitrogen flux. After 5 min of reaction, 1 mL of N,N-dimethylformamide (DMF) solvent was added to keep an efficient stirring. This solvent addition was repeated after 60 min of reaction. After 4 h of reaction, a determined amount of 1,4-butanediol (1,4-BDO) (20 mg) was added and the polymerization was allowed to proceed for an additional 1 h. The reaction product was then dried under vacuum at 80 °C for 16 hours to remove DMF. The final product was recovered as a yellowish solid.

### 4.9 Analytics

#### 4.9.1 PHA extraction and content determination

The polymer content was assayed by subjecting the lyophilized cells to acidic methanolysis as previously described^69^. The PHA monomers’ methylesters were assayed by GC using a Hewlett-Packard 6890 N chromatograph equipped with a HP-Innowax capillary column (30 m × 0.25 mm, 0.50-μm film thickness; Agilent Technologies) and a flame ionization detector (FID), using the temperature program previously described^70^. Total PHA content was determined as a percentage of cell dry weight (CDW).

#### 4.9.2 Nitrogen quantification

The concentration of nitrogen in the media was monitored by taking 1 ml samples from the cultures at various time points and centrifuging them for 3 min at 16900 × *g* (benchtop 5430R centrifuge; Eppendorf, Germany). The supernatant was retained and the nitrogen concentration was determined using the method of Scheiner^71^. Briefly, the supernatant was diluted to 10^−3^ in deionized water and placed in a 1 cm path length, 1.6 ml volume cuvette (Sarstedt, Germany). 400 µl of phenol solution (per 100 ml of deionized water: 1.3 g Na_3_PO_4_; 3 g Na_3_C_3_H_5_O_7_; 0.3 g sodium EDTA; 6 g phenol; 0.02 g sodium nitroprusside) was added to the cuvette and mixed well. This was promptly followed by the addition of 600 µl of alkaline solution (per 100 ml of final volume: 40 ml 1 M NaOH; 2.5 ml hypochlorite solution; 57.5 ml deionized water). The mixture was incubated at room temperature in the dark for 45 min. The formation of indophenol-blue was measured at 635 nm using the Unicam Helios δ UV/VIS spectrophotometer (Thomas Scientific, USA).

#### 4.9.3 HAA quantification

HAA was quantified using high-performance liquid-chromatography coupled with a charged aerosol detector using a method established earlier^47^.

#### 4.9.4 EG quantification

At UCD, EG depletion was monitored using on an Aminex HPX-87H ion exclusion column (300 mm x 7.8 mm, particle size 9 μm; Bio-rad). The column was maintained at 40°C and samples were isocratically eluted using 0.014 N H_2_SO_4_ at a flow rate of 0.55 ml min^-1^ and read on a refractive index detector (RID). The EG retention time under the above conditions was 23 min.

In the iAMB labs, an ion exchange chromatography was applied for EG quantification. The used System Gold was composed of a pump LC-126, an autosampler LC-508, a UV detector LC-166, (all Beckmann Coulter, Krefeld, Germany), a Jetstream 2 Plus column oven (Knauer, Berlin, Germany), and a refractive index detector Smartline RI Detector 2300 (Knauer, GmbH, Berlin, Germany). The applied column was the Metab-AAC (ISERA GmbH, Düren, Germany) with a length of 30 cm and a diameter of 7.8 mm. The running buffer was 5 mM sulfuric acid, which was pumped isocratically with a flow rate of 0.8 mL/min at a temperature of 80 °C. 20 µL of the sample were injected.

#### 4.9.5 TA quantification

In the UCD lab, the supernatant collected during growth was diluted 20-fold and filtered (500 μl) using Mini-UniPrep syringeless filter devices (GE Healthcare Life Science, Ireland). TA concentration was analyzed according to the protocol previously outlined by Kenny^72^. At the iAMB, the amounts of the UV-absorbing degradation products in the STR samples were measured by reversed-phase HPLC as described before^33^. A C18 column (Eurospher 100-5, 150 mm × 4.6 mm with pre-column, Knauer GmbH, Berlin, Germany) and a mobile phase consisting of 20% acetonitrile, 20% 10 mM sulfuric acid and 60% ultra-pure water was used. The hydrolysis supernatant samples were diluted using the mobile phase, acidified with concentrated HCl (37%) and then centrifuged to remove any precipitation. The detection of TA and associated low-molecular-weight (LMW) esters was performed at a wavelength of 241 nm.

#### 4.9.6 HAA-based poly(amide urethane) characterization

^1^H- and ^13^C-NMR spectra were obtained with a Bruker 400 MHz spectrophotometer. CDCl_3_ and DMSO-d_6_ were used as deuterated solvent to prepare solutions with concentrations of 8-10 and 30-50 mg/mL for ^1^H-NMR and ^13^C-NMR, respectively. The number of scans was set to 128 and 1024 for ^1^H- and ^13^C-NMR, respectively. Spectra were calibrated using the CDCl_3_ peak (δ_H_ = 7.26 ppm, δ_C_ = 77.16 ppm) or the DMSO-d_6_ peak (δ_H_ = 2.50 ppm, δ_C_ = 39.52 ppm).

Fourier transformed infrared spectroscopy (FTIR) was performed with a Nicolet 380 spectrometer (Thermo Electron Corporation) used in reflection mode and equipped with an ATR diamond module (FTIR-ATR). The FTIR-ATR spectra were collected at a resolution of 4 cm^-1^ and with 32 scans per run.

Differential scanning calorimetry (DSC) was performed using a TA Instrument Q200. Samples of 2-3 mg in sealed aluminum pans were analyzed under nitrogen flow (50 mL/min). A three-step procedure with a 10 °C/min ramp was applied as follow: (1) heating up from room temperature to 200 °C and holding for 3 min to erase the thermal history, (2) cooling down to −60 °C and holding for 3 min, (3) heating up (second heating) from −60 °C to 200 °C.

Thermal stability was studied by thermogravimetric analyses (TGA). Measurements were conducted under air atmosphere (flow rate of 25 mL/min) using a Hi-Res TGA Q5000 apparatus from TA Instruments. Samples (1-3 mg) were heated from room temperature up to 800 °C at a rate of 10 °C/min.

### 4.10 Flux balance analysis

FBA has been carried out as described previously^73^. Briefly, the genome-scale model of *P. putida* KT2440, *i*JN1411, was used^74^ and extended by the biosynthesis routes for HAA production and the metabolization routes for EG and TA. All simulations were carried out in MATLAB (version R2017b, The Mathworks, Inc., Natick, MA, USA) using the COBRA toolbox^75^, with the linear programming solver of Gurobi (www.gurobi.com).

## Abbreviations

EG: ethylene glycol,
TA: terephthalic acid terephthalate,
PET: polyethylene terephthalate,
PHA: polyhydroxyalkanoate,
HAA: hydroxyalkanoyloxy-alkanoate,
MHET: mono-(2-hydroxyethyl)TA

## 5 Acknowledgements

The authors have received funding from the European Union’s Horizon 2020 research and innovation program under grant agreement no. 633962 for the project P4SB. TN is funded by Science Foundation Ireland grant number 16/RC/3889.

TT and LMB have been partially funded by the Deutsche Forschungsgemeinschaft (DFG, German Research Foundation) under Germany’s Excellence Strategy – Exzellenzcluster 2186

“The Fuel Science Center” ID: 390919832.

## 6 Author Contributions

TT supervised the experiments regarding monomer metabolism and HAA synthesis, drafted the manuscript, and coordinated the study, TN provided strain *Pseudomonas* sp. GO16, supervised the experiments regarding PHA synthesis and drafted parts of the manuscript, RW supervised the experiments regarding depolymerization and drafted parts of the manuscript, EP supervised the experiments regarding polymerization and drafted parts of the manuscript, KS carried out the experiments regarding monomer metabolism and HAA synthesis, NB carried out the experiments regarding PHA synthesis, AH carried out the experiments regarding depolymerization, MJ carried out the experiments regarding polymerization, SK was involved in PHA bioprocess design, NW was involved in designing and coordinating the study, drafted parts of the manuscript and critically read the manuscript, RP was involved in designing the study and critically read the manuscript, LA was involved in designing the study and critically read the manuscript, WZ was involved in designing the study and critically read the manuscript, KOC designed the study and critically read the manuscript, LMB designed and coordinated the study and critically read the manuscript. All authors read and approved the final manuscript.

