## Supplementary information for "Bio-upcycling of polyethylene terephthalate"

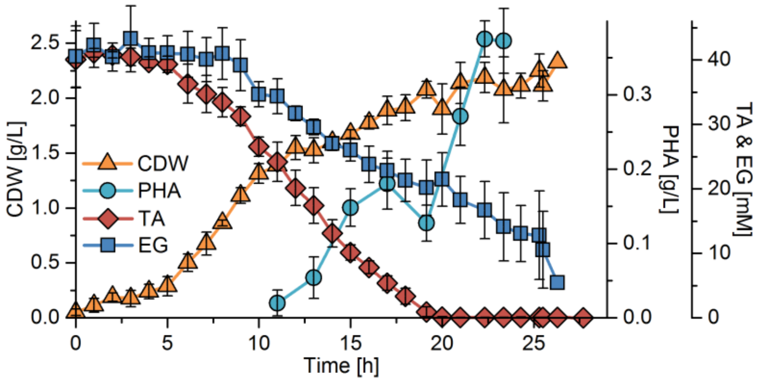


**Fig. S1** Growth, PHA accumulation, and substrate depletion by *Pseudomonas* sp. GO16 KS3 when a synthetic mixture of TA and EG was used under nitrogen limiting conditions. *Pseudomonas* sp. GO16 KS3 was cultivated in a 5 L bioreactor with 3 L of mineral salts medium (MSM) at 30 °C. The PET monomers were added to a concentration of 40 mM TA and 40 mM EG. Growth (cell dry weight, CDW), polyhydroxyalkanoate (PHA, %CDW), substrates terephthalic acid (TA) and ethylene glycol (EG). The error bars represent the standard deviation from the mean of three independent biological replicates.
